# Library preparation method and DNA source influence endogenous DNA recovery from 100-year-old avian museum specimens

**DOI:** 10.1101/2023.02.06.527363

**Authors:** Amie E. Settlecowski, Ben D. Marks, Joseph D. Manthey

## Abstract

Museum specimens collected prior to cryogenic tissue storage are increasingly being used as genetic resources, and though high throughput sequencing is becoming more cost-efficient, whole genome sequencing (WGS) of historical DNA (hDNA) remains inefficient and costly due to its short fragment sizes and high loads of exogenous DNA, among other factors. It is also unclear how sequencing efficiency is influenced by DNA source. We aimed to identify the most efficient method and DNA source for collecting WGS data from avian museum specimens. We analyzed low-coverage WGS from 60 DNA libraries prepared from four American Robin (*Turdus migratorius*) and four Abyssinian Thrush (*Turdus abyssinicus*) specimens collected in the 1920s. We compared DNA source (toepad versus incision-line skin clip) and three library preparation methods: 1) double-stranded, single tube (KAPA); 2) single-stranded, multi-tube (IDT); and 3) single-stranded, single-tube (Claret Bioscience). We found that the multi-tube ssDNA method resulted in significantly greater endogenous DNA content, average read length, and sequencing efficiency than the other tested methods. We also tested whether a predigestion step reduced exogenous DNA in libraries from one specimen per species and found promising results that warrant further study. The ~10% increase in average sequencing efficiency of the best performing method over a commonly implemented dsDNA library preparation method has the potential to significantly increase WGS coverage of hDNA from bird specimens. Future work should evaluate the threshold for specimen age at which these results hold and how the combination of library preparation method and DNA source influence WGS in other taxa.

## Introduction

Museum specimens collected prior to cryogenic tissue storage have long been used as genetic resources to address questions in ecology, evolutionary biology, and conservation (Habel, Husemann, Finger, Danley, & Zachos, 2014; Wandeler, Hoeck, & Keller, 2007). Genetic studies using these specimens have increased with the advent of high throughput sequencing methods, which in comparison to prior Sanger sequencing methods, drastically increased the data returned from each destructive sampling (Burrell, Disotell, & Bergey, 2015). Now museum specimens commonly facilitate genomic studies via reduced representation (Bi et al., 2013; Linck, Hanna, Sellas, & Dumbacher, 2017; McCormack, Tsai, & Faircloth, 2016) and even whole genome sequencing (e.g. van der Valk, Díez-del-Molino, Marques-Bonet, Guschanski, & Dalén, 2019; Wu et al., 2022). Despite its increasing prevalence and dropping cost, sequencing whole genomes of museum specimens remains expensive because of the degraded nature of the historical DNA (hDNA).

Historical DNA tends to consist of short fragment lengths (McDonough, Parker, McInerney, Campana, & Maldonado, 2018; Straube et al., 2021; Tsai, Schedl, Maley, & McCormack, 2020) that are smaller than the recommended library sizes for the most cost-efficient sequencing setups (i.e., Illumina NovaSeq 6000 S4 flowcell, 200 or 300 cycles). As a result, many sequencing cycles are directly wasted by a lack of base pairs to sequence or indirectly wasted on adapter read through (Straube et al., 2021). Historical DNA libraries also tend to consist of low proportions of DNA from the focal specimen (hereafter endogenous DNA). The rest of the library may consist of exogenous DNA from (1) microbes that have colonized the museum specimen or (2) other environmental microbes, (3) contaminating DNA from researchers or (4) other museum specimens, and (5) more recent DNA samples (Fulton & Shapiro, 2019). Altogether, the degraded nature of hDNA results in the recovery of lower proportions of endogenous DNA sequence data (Burrell et al., 2015) and necessitates increased sequencing effort per specimen to recover similar WGS coverage to modern, high quality DNA libraries. This inefficiency limits the use of historical DNA from museum specimens to address population genomic questions that require larger sample sizes in addition to sufficient coverage to address questions about selection and demography (Lou, Jacobs, Wilder, & Therkildsen, 2021).

Ancient DNA researchers have identified that single-stranded (Gansauge & Meyer, 2013) and single-tube library preparation methods (Carøe et al., 2018), and those that ligate adapters to unmodified DNA molecule ends (Kapp, Green, & Shapiro, 2021), increase the amount of degraded ancient DNA molecules that are converted into genomic libraries. However, the most implemented ancient library preparation methods are non-proprietary (Gansauge et al., 2017; Henneberger, Barlow, & Paijmans, 2019), thus requiring a high level of startup effort. Early ssDNA methods were also more expensive to implement than double-stranded DNA (dsDNA) libraries, and their improvement in sequencing efficiency did not warrant the additional effort and cost to implement for all but the most degraded ancient DNA samples (Wales et al., 2015). That is perhaps why only two studies to date have evaluated the influence of ssDNA versus dsDNA library preparation on shotgun sequencing of historical specimens. Sproul and Maddison (2017) found that ssDNA libraries–in comparison to dsDNA libraries–prepared from 16 whole beetle specimens resulted in more retained reads following quality filtering, but no difference in endogenous DNA content. Similarly, Hahn et al. (2022) recently found no difference in endogenous DNA content or insert length between ssDNA and dsDNA libraries prepared from twelve taxonomically diverse wet collection vertebrate specimens. Additional studies of the influence of library preparation on WGS of museum specimens that control for taxonomy, locality, and collection age of specimens will be valuable moving forward. Early ssDNA methods have been modified to reduce costs and ease implementation (Gansauge et al., 2017) and ssDNA methods are now commercially available as kits facilitating further study of their impact on hDNA sequencing efficiency.

Thus far, the majority of research investigating how to maximize the recovery of genetic data from non-ancient museum specimens has focused on the influence of DNA source or extraction method on DNA yield (Hahn et al., 2022; Hawkins, Flores, McGowen, & Hinckley, 2022; McDonough et al., 2018; Pacheco et al., 2022; Straube et al., 2021; Tsai et al., 2020; Zacho et al., 2021). However, DNA yield does not necessarily predict sequencing success or efficiency (McDonough et al., 2018; Straube et al., 2021; Zacho et al., 2021) because it is not possible to estimate the proportions of endogenous versus exogenous extracted DNA. For example, a recent study of hundreds of historical genomic DNA libraries built from samples of museum bird specimens found that those built from specimens of smaller species (which generally produce smaller samples) unintuitively had a higher proportion of endogenous sequence data (Irestedt et al., 2022). Moreover, a few studies have shotgun sequenced DNA from multiple sources on the same specimen and found differences in endogenous DNA content across sampling sites in fluid-preserved garter snake specimens (Zacho et al., 2021), prepared mammal skins (McDonough et al., 2018), and formalin-fixed specimens of a dozen vertebrate taxa (Hahn et al., 2022). Despite research indicating that differences in hDNA sourced from toepads versus incision-line clips in bird specimens could influence high-throughput sequencing results, no studies have evaluated their difference in endogenous DNA content and sequencing efficiency.

Bird study skin specimens have been an especially prolific source of hDNA research (Billerman & Walsh, 2019) in part due to preservation methods (i.e., skin drying) that are not catastrophic to DNA preservation relative to methods such as formalin-fixation. Bird study skins have been the foci of some of the earliest studies of hDNA (Mundy, Unitt, & Woodruff, 1997), the source for some of the first implementations of reduced representation, high throughput sequencing methods using museum specimens (Linck et al., 2017; McCormack et al., 2016), and some of the largest studies implementing WGS of hDNA to date (Irestedt et al., 2022). In this study we aim to maximize the potential of hDNA from bird study skins by identifying whether DNA source and library preparation method influence the endogenous DNA content and sequencing efficiency of hDNA libraries, and by introducing a pre-digestion step prior to DNA extraction to reduce exogenous DNA.

In this study we test three library preparation methods that vary in 1) the number of cleanups and tube transfers that occur before the library amplification step (one vs two) and 2) whether they convert single or double-stranded DNA into library molecules. Each cleanup and tube transfer is an opportunity to lose DNA molecules of the target length (greater than number of sequencing cycles) due to the inherent imprecision of SPRI bead cleanups in addition to pipette error. Methods that are optimized with one, in comparison to two, tube transfers should transform more DNA molecules of target length into library molecules, thus maximizing library complexity and sequencing efficiency. Double-stranded DNA library preparation methods cannot convert ssDNA into library molecules, though as described above, hDNA is expected to consist in some proportion of single strand molecules due to degradation. Thus, we expect that dsDNA libraries prepared from hDNA will have reduced sequencing efficiency and possibly endogenous DNA content compared to that of ssDNA libraries.

We also test the influence of DNA source—toepads versus incision-line skin clips—on endogenous DNA content and sequencing efficiency. A previous study indicated that toepads consist of longer DNA fragments than skin clips (Tsai et al., 2020), another possible source of hDNA from birds (Töpfer, Gamauf, & Haring, 2011). Libraries prepared from samples consisting of longer DNA fragments should maximize the sequencing capacity, resulting in longer read lengths on average and greater sequencing efficiency. To test these expectations, we prepared shotgun DNA libraries from a toepad and skin clip from eight approximately 100-year-old bird specimens via three methods: 1) double-stranded, single tube (KAPA HyperPrep Kit); 2) single-stranded, multi-tube (IDT xGen™ ssDNA & Low-Input DNA Library Prep Kit); and 3) single-stranded, single-tube (Claret Bioscience SRSLY® NanoPlus Kit). We sought to reduce exogenous DNA by implementing a predigestion step prior to DNA extraction, to our knowledge, for the first time on bird specimens. To qualitatively evaluate the influence of the predigestion we also prepared libraries from replicate toepad and skin clip DNA extractions not subjected to predigestion from two of the eight specimens (Figure 1).

**Figure 1.**
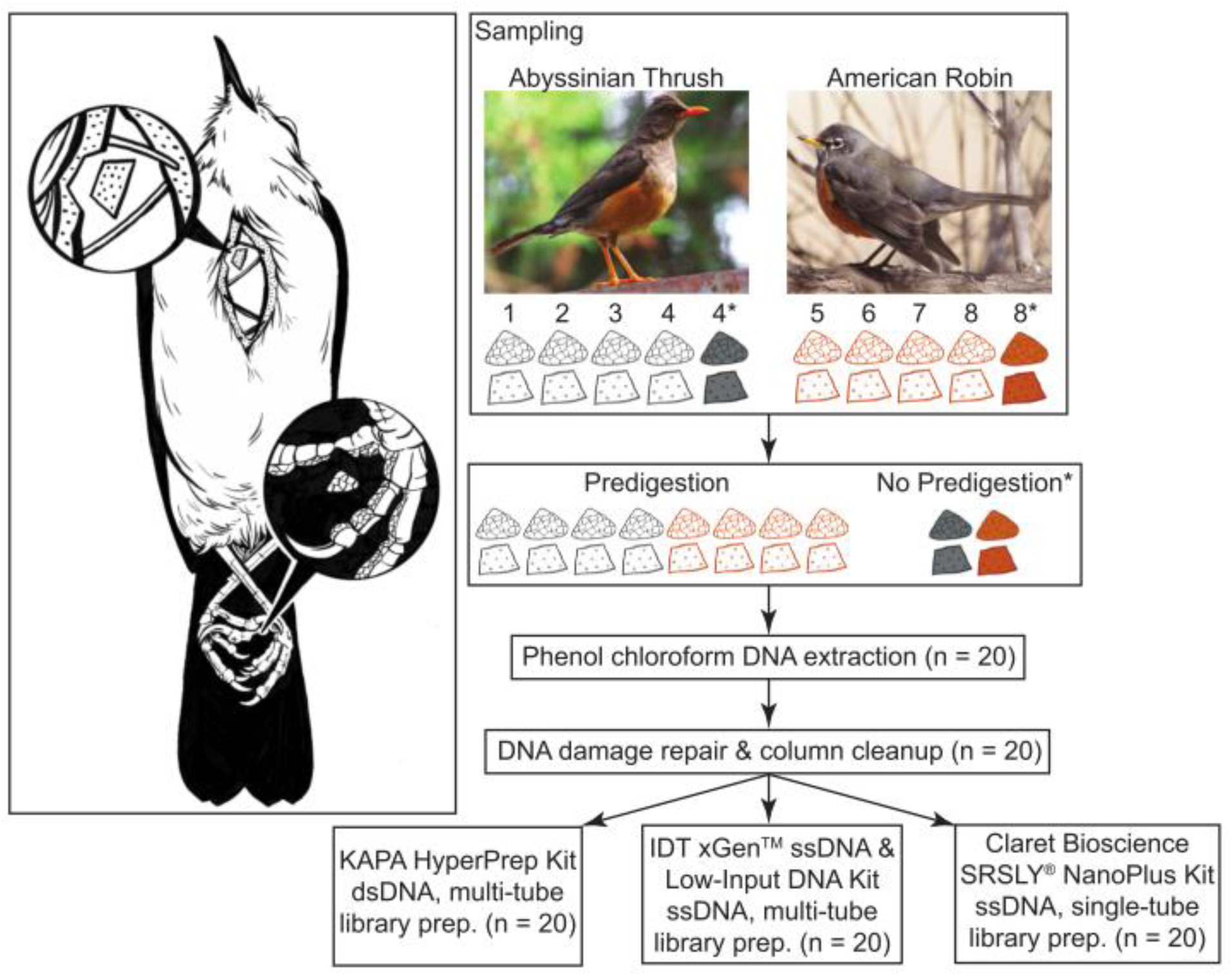
Study sampling design. The illustration on the left shows where incision-line skin clips and toepads are sampled from the specimens. The flowchart on the right is an overview of the experimental design and the laboratory procedures. Toepads and skin clips are depicted as triangular and trapezoidal shapes, respectively. Photographs by JDM and illustrations by L. Nassef.

## METHODS

### 2.1. Sampling

We sampled eight bird specimens: four Abyssinian Thrush (*Turdus abyssinicus*; hereafter thrushes) and four American Robin (*Turdus migratorius*; hereafter robins; Table 1). We chose the thrush specimens based on their inclusion in another ongoing project and chose to bolster our sample size for this study with the robin specimens because they are a closely related, similar species that is well-represented in North American natural history collections. Moreover, the thrushes were collected in the tropics and the robins were collected in a temperate region which could influence the drying time of the study skins and in turn, possible degradation due to rot or the microbial load within dried skin (Irestedt et al., 2022). We chose specimens collected within one year of each other to control for DNA degradation due to time since specimen preparation.

**Table 1.** Summary of extracted DNA and sequencing results for each sample.

| Species | Sample | Post-extraction |  | Post-repair |  | Method | Type | Raw reads<br>(millions) | Clean reads<br>(millions) | Mapped<br>reads<br>(millions) | Mean<br>read<br>length | Dist.to<br>ref. | Endog.<br>DNA<br>content | Seq.<br>eff. |
| --- | --- | --- | --- | --- | --- | --- | --- | --- | --- | --- | --- | --- | --- | --- |
|  |  | Yield<br>(ng/mg) | Mean<br>DNA<br>size (bp) | Yield<br>(ng/mg) | Mean<br>DNA<br>size (bp) |  |  |  |  |  |  |  |  |  |
| *T. abyssinicus* | 83107 |  |  |  |  | IDT | skin | 10.54 | 3.38 | 2.96 | 71.99 | 0.018 | 0.851 | 0.193 |
| *T. abyssinicus* | 83107 | 400.03 | 57.36 | 136.10 | 68.02 | KAPA | skin | 60.29 | 15.17 | 12.45 | 59.75 | 0.014 | 0.820 | 0.119 |
| *T. abyssinicus* | 83107 |  |  |  |  | SRSLY | skin | 12.85 | 5.23 | 4.64 | 75.14 | 0.015 | 0.822 | 0.267 |
| *T. abyssinicus* | 83107 |  |  |  |  | IDT | toepad | 12.62 | 5.51 | 5.12 | 97.84 | 0.017 | 0.900 | 0.382 |
| *T. abyssinicus* | 83107 | 698.97 | 65.75 | 332.89 | 83.90 | KAPA | toepad | 85.06 | 28.05 | 25.05 | 78.48 | 0.012 | 0.889 | 0.224 |
| *T. abyssinicus* | 83107 |  |  |  |  | SRSLY | toepad | 8.81 | 3.36 | 2.89 | 67.02 | 0.017 | 0.853 | 0.213 |
| *T. abyssinicus* | 83109 |  |  |  |  | IDT | skin | 11.26 | 3.29 | 1.69 | 62.71 | 0.035 | 0.480 | 0.086 |
| *T. abyssinicus* | 83109 | 376.78 | 57.93 | 73.28 | 61.74 | KAPA | skin | 50.69 | 12.09 | 5.77 | 53.49 | 0.027 | 0.477 | 0.057 |
| *T. abyssinicus* | 83109 |  |  |  |  | SRSLY | skin | 10.08 | 3.74 | 2.13 | 65.56 | 0.030 | 0.519 | 0.119 |
| *T. abyssinicus* | 83109 |  |  |  |  | IDT | toepad | 10.95 | 4.56 | 4.18 | 88.02 | 0.018 | 0.896 | 0.323 |
| *T. abyssinicus* | 83109 | 858.85 | 66.68 | 338.13 | 85.08 | KAPA | toepad | 63.05 | 21.06 | 18.86 | 79.73 | 0.013 | 0.890 | 0.232 |
| *T. abyssinicus* | 83109 |  |  |  |  | SRSLY | toepad | 9.29 | 3.46 | 2.97 | 66.34 | 0.017 | 0.853 | 0.206 |
| *T. abyssinicus* | 83114 |  |  |  |  | IDT | skin | 46.55 | 32.78 | 30.90 | 100.17 | 0.009 | 0.916 | 0.640 |
| *T. abyssinicus* | 83114 | 1350.36 | 78.71 | 576.64 | 218.79 | KAPA | skin | 32.03 | 15.21 | 14.27 | 109.38 | 0.013 | 0.919 | 0.472 |
| *T. abyssinicus* | 83114 |  |  |  |  | SRSLY | skin | 9.56 | 5.06 | 4.69 | 104.24 | 0.017 | 0.903 | 0.503 |
| *T. abyssinicus* | 83114 |  |  |  |  | IDT | toepad | 24.48 | 10.02 | 8.80 | 89.44 | 0.016 | 0.856 | 0.309 |
| *T. abyssinicus* | 83114 | 800.00 | 53.96 | 55.17 | 98.74 | KAPA | toepad | 45.38 | 15.44 | 13.93 | 83.94 | 0.015 | 0.889 | 0.250 |
| *T. abyssinicus* | 83114 |  |  |  |  | SRSLY | toepad | 7.62 | 3.34 | 3.01 | 85.32 | 0.018 | 0.874 | 0.332 |
| *T. abyssinicus* | 83114<sup>†</sup> |  |  |  |  | IDT | skin | 42.34 | 28.30 | 26.73 | 101.24 | 0.009 | 0.917 | 0.615 |
| *T. abyssinicus* | 83114<sup>†</sup> | 4254.55 | 96.90 | 1872.73 | 211.98 | KAPA | skin | 34.26 | 15.74 | 14.75 | 107.05 | 0.013 | 0.919 | 0.447 |
| *T. abyssinicus* | 83114<sup>†</sup> |  |  |  |  | SRSLY | skin | 9.35 | 4.80 | 4.44 | 101.67 | 0.017 | 0.902 | 0.474 |
| *T. abyssinicus* | 83114<sup>†</sup> |  |  |  |  | IDT | toepad | 28.09 | 11.70 | 10.77 | 99.56 | 0.015 | 0.891 | 0.368 |
| *T. abyssinicus* | 83114<sup>†</sup> | 1901.96 | 56.07 | 202.94 | 107.53 | KAPA | toepad | 62.51 | 22.23 | 20.22 | 84.52 | 0.013 | 0.899 | 0.266 |
| *T. abyssinicus* | 83114<sup>†</sup> |  |  |  |  | SRSLY | toepad | 14.23 | 6.31 | 5.75 | 85.21 | 0.016 | 0.876 | 0.344 |
|  |  | Yield<br>(ng/mg) | Mean<br>DNA<br>size (bp) | Yield<br>(ng/mg) | Mean<br>DNA<br>size (bp) |  |  |  |  |  |  |  |  |  |
| *T. abyssinicus* | 83115 |  |  |  |  | IDT | skin | 14.52 | 4.28 | 3.80 | 78.74 | 0.017 | 0.858 | 0.197 |
| *T. abyssinicus* | 83115 | 942.98 | 56.71 | 526.94 | 69.01 | KAPA | skin | 80.02 | 19.29 | 16.20 | 59.36 | 0.014 | 0.843 | 0.117 |
| *T. abyssinicus* | 83115 |  |  |  |  | SRSLY | skin | 11.55 | 3.30 | 2.65 | 54.80 | 0.015 | 0.803 | 0.120 |
| *T. abyssinicus* | 83115 |  |  |  |  | IDT | toepad | 43.02 | 17.60 | 16.22 | 94.73 | 0.013 | 0.893 | 0.345 |
| *T. abyssinicus* | 83115 | 564.00 | 59.43 | 217.47 | 83.13 | KAPA | toepad | 14.25 | 4.92 | 4.37 | 78.08 | 0.018 | 0.881 | 0.232 |
| *T. abyssinicus* | 83115 |  |  |  |  | SRSLY | toepad | 12.70 | 5.55 | 5.02 | 81.83 | 0.015 | 0.862 | 0.324 |
| *T. migratorius* | 162188 |  |  |  |  | IDT | skin | 15.77 | 4.77 | 3.97 | 60.97 | 0.018 | 0.822 | 0.146 |
| *T. migratorius* | 162188 | 375.19 | 52.58 | 597.41 | 58.71 | KAPA | skin | 2.14 | 0.74 | 0.60 | 60.52 | 0.018 | 0.803 | 0.162 |
| *T. migratorius* | 162188 |  |  |  |  | SRSLY | skin | 9.77 | 3.52 | 2.94 | 61.09 | 0.018 | 0.823 | 0.176 |
| *T. migratorius* | 162188 |  |  |  |  | IDT | toepad | 21.99 | 9.50 | 8.58 | 95.19 | 0.016 | 0.879 | 0.354 |
| *T. migratorius* | 162188 | 501.80 | 57.60 | 436.09 | 76.70 | KAPA | toepad | 10.85 | 4.53 | 3.96 | 78.38 | 0.019 | 0.866 | 0.275 |
| *T. migratorius* | 162188 |  |  |  |  | SRSLY | toepad | 8.36 | 3.60 | 3.15 | 76.19 | 0.019 | 0.868 | 0.279 |
| *T. migratorius* | 162188<sup>†</sup> |  |  |  |  | IDT | skin | 6.56 | 4.11 | 0.86 | 80.05 | 0.020 | 0.174 | 0.100 |
| *T. migratorius* | 162188<sup>†</sup> | 173.57 | 54.16 | 47.37 | 62.90 | KAPA | skin | 14.98 | 6.38 | 2.90 | 59.05 | 0.018 | 0.334 | 0.110 |
| *T. migratorius* | 162188<sup>†</sup> |  |  |  |  | SRSLY | skin | 8.85 | 3.79 | 2.07 | 63.26 | 0.017 | 0.415 | 0.135 |
| *T. migratorius* | 162188<sup>†</sup> |  |  |  |  | IDT | toepad | 9.14 | 3.95 | 3.55 | 89.65 | 0.019 | 0.875 | 0.333 |
| *T. migratorius* | 162188<sup>†</sup> | 231.03 | 59.13 | 98.59 | 77.89 | KAPA | toepad | 31.17 | 12.31 | 10.86 | 78.33 | 0.017 | 0.873 | 0.263 |
| *T. migratorius* | 162188<sup>†</sup> |  |  |  |  | SRSLY | toepad | 10.20 | 4.42 | 3.92 | 78.11 | 0.019 | 0.870 | 0.293 |
| *T. migratorius* | 175413 |  |  |  |  | IDT | skin | 8.18 | 2.72 | 2.31 | 78.30 | 0.019 | 0.807 | 0.208 |
| *T. migratorius* | 175413 | 197.25 | 50.97 | 59.33 | 62.37 | KAPA | skin | 15.67 | 4.99 | 3.86 | 60.91 | 0.018 | 0.770 | 0.144 |
| *T. migratorius* | 175413 |  |  |  |  | SRSLY | skin | 11.30 | 4.57 | 3.84 | 70.22 | 0.017 | 0.792 | 0.227 |
| *T. migratorius* | 175413 |  |  |  |  | IDT | toepad | 12.34 | 6.03 | 5.66 | 100.66 | 0.017 | 0.894 | 0.442 |
| *T. migratorius* | 175413 | 501.09 | 56.22 | 219.23 | 86.23 | KAPA | toepad | 32.87 | 13.07 | 11.71 | 81.68 | 0.016 | 0.882 | 0.280 |
| *T. migratorius* | 175413 |  |  |  |  | SRSLY | toepad | 7.51 | 3.53 | 3.24 | 86.20 | 0.017 | 0.868 | 0.372 |
|  |  | Yield<br>(ng/mg) | Mean<br>DNA<br>size (bp) | Yield<br>(ng/mg) | Mean<br>DNA<br>size (bp) |  |  |  |  |  |  |  |  |  |
| *T. migratorius* | 175421 |  |  |  |  | IDT | skin | 6.68 | 2.31 | 1.97 | 64.10 | 0.019 | 0.840 | 0.180 |
| *T. migratorius* | 175421 | 463.63 | 52.32 | 111.02 | 62.95 | KAPA | skin | 56.18 | 17.57 | 14.70 | 63.39 | 0.015 | 0.833 | 0.159 |
| *T. migratorius* | 175421 |  |  |  |  | SRSLY | skin | 14.82 | 5.83 | 5.05 | 67.82 | 0.017 | 0.838 | 0.222 |
| *T. migratorius* | 175421 |  |  |  |  | IDT | toepad | 21.44 | 9.64 | 8.98 | 98.88 | 0.016 | 0.891 | 0.398 |
| *T. migratorius* | 175421 | 838.10 | 60.18 | 234.00 | 82.44 | KAPA | toepad | 51.02 | 19.65 | 17.51 | 78.30 | 0.015 | 0.879 | 0.259 |
| *T. migratorius* | 175421 |  |  |  |  | SRSLY | toepad | 7.29 | 3.33 | 3.03 | 83.03 | 0.018 | 0.868 | 0.343 |
| *T. migratorius* | 66823 |  |  |  |  | IDT | skin | 60.98 | 51.78 | 49.16 | 95.45 | 0.007 | 0.920 | 0.736 |
| *T. migratorius* | 66823 | 1150.86 | 387.24 | 950.45 | 1923.43 | KAPA | skin | 35.56 | 24.12 | 22.91 | 106.55 | 0.010 | 0.921 | 0.659 |
| *T. migratorius* | 66823 |  |  |  |  | SRSLY | skin | 9.88 | 6.23 | 5.87 | 105.58 | 0.017 | 0.917 | 0.605 |
| *T. migratorius* | 66823 |  |  |  |  | IDT | toepad | 5.13 | 2.24 | 2.03 | 85.15 | 0.020 | 0.885 | 0.321 |
| *T. migratorius* | 66823 | 223.70 | 53.26 | 47.36 | 75.28 | KAPA | toepad | 50.77 | 19.75 | 17.26 | 73.40 | 0.015 | 0.871 | 0.241 |
| *T. migratorius* | 66823 |  |  |  |  | SRSLY | toepad | 11.86 | 4.31 | 3.63 | 61.16 | 0.018 | 0.855 | 0.182 |

†Extraction replicate not subjected to predigestion

We collected two samples from each specimen—a toepad and a “skin punch” from the incision-line through the pectoral apterium (following Tsai et al., 2020)—to evaluate whether tissue source differed in proportion of endogenous DNA. We also took replicate samples from one specimen of each species (Table 1) to qualitatively evaluate the effect of sample predigestion prior to DNA extraction on the exogenous DNA load. The experimental design is summarized in Figure 1.

We followed stringent sampling precautions to limit the introduction of contaminant DNA: We (1) wore surgical masks and gloves throughout sampling, (2) took samples in the collections away from any specimen preparation laboratory, (3) did not enter any modern molecular DNA or specimen preparation laboratory prior to sampling, (4) prepared the work surface and all other supplies (e.g., forceps, optivisor, writing utensil) by cleaning with freshly prepared 10% bleach followed by 70% isopropanol or ethanol, and (5) immediately placed samples in sterile microcentrifuge tubes that were unpackaged in a sterile lab and not opened prior to beginning molecular lab work. We used a fresh pair of gloves and sterile scalpel blade for each sample to minimize contamination between samples.

### 2.2. Molecular laboratory work

We followed stringent ancient DNA clean lab protocols to minimize contamination during molecular laboratory work (Fulton & Shapiro, 2019). We completed all pre-PCR steps in an ancient DNA facility in the Department of Human Genetics at the University of Chicago in a non-human specific room. We performed each step prior to PCR in a maximum batch size of 12 samples, introduced a negative control in each batch of extractions and library preparations, and then carried these controls through the remaining steps of lab work.

We extracted DNA via a phenol-chloroform protocol followed by ethanol precipitation with minor modifications to the protocol presented in Tsai et al. (2020). We performed an NEB PreCR DNA repair treatment following the sequential reaction protocol on each DNA extraction. This treatment repairs DNA damage from hydrolysis and oxidative stress, among other mechanisms, that results in deaminated cytosines, nicks, and other DNA damage incurred with age. A previous study found that a different NEB repair kit optimized for formalin-fixed specimens increased the yields of libraries prepared from historical beetle specimens by approximately 30% (Sproul & Maddison, 2017). Following the damage repair treatment, we performed a Qiagen MinElute column cleanup and resuspended the DNA in 50 μL of PCR-grade water. Next, we measured the DNA yield and distribution of DNA fragment sizes using the Qubit High Sensitivity dsDNA (Thermo Fisher Scientific) and Agilent Bioanalyzer High-Sensitivity DNA Kit assays, respectively, following the DNA extraction and again following DNA repair and cleanup. We performed the same assays for each extraction negative control to monitor for contamination.

We prepared three shotgun sequencing libraries from each DNA extraction and negative control. Each of the three libraries were prepared via a different method: 1) double-stranded, single tube (KAPA HyperPrep Kit); 2) single-stranded, multi-tube (IDT xGen™ ssDNA & Low-Input DNA Library Prep Kit); and 3) single-stranded, single-tube (Claret Bioscience SRSLY® NanoPlus Kit). We largely followed manufacturer protocols with the following modifications: during the KAPA adapter ligation step we ligated 25 μM iTru Stubs (Glenn et al., 2019) to each library molecule. For all cleanups we used a homebrew SPRI bead-solution (Rohland & Reich, 2012) and for each cleanup step in the KAPA and IDT protocols we performed 1.2x SPRI concentration cleanups. We indexed each library via amplification with 2.5 μM of a unique pair of iTru5 and iTru7 indexed primers (Glenn et al., 2019) and KAPA HiFi HotStart Uracil+ ReadyMix. For amplification we split each adapter-ligated library into two replicates of 25 μL each and ran a nine- to twelve-cycle PCR, depending on input DNA amount and method, on the first replicate; then we estimated the yield of the first replicate via a Qubit High Sensitivity dsDNA assay and ran a 10- to 12-cycle PCR on the second adapter-ligated library replicate. We combined amplified replicates for each library and performed a final SPRI cleanup. Finally, we measured the average molecule size and calculated the concentration of adapter ligated molecules for each sample library via an Agilent Bioanalyzer High-Sensitivity DNA Kit assay and qPCR with the KAPA Library Quantification Kit. We submitted a final library pool to Texas Tech University Center for Biotechnology & Genomics for sequencing. They first checked that libraries were sequencable with an Illumina MiSeq nano run followed by 100 base pair (bp) paired-end sequencing on one Illumina NovaSeq SP flowcell.

### 2.3. Bioinformatics

We received demultiplexed sequence data as raw fastq files from the sequencing facility and ran FastQC (https://www.bioinformatics.babraham.ac.uk/projects/fastqc/) to assess quality and adapter contamination by library preparation method. We trimmed 10 bp from the beginning of every IDT library read 2 via Seqtk *trimfq* (https://github.com/lh3/seqtk) to remove a low-complexity polynucleotide tail that facilitates adapter ligation in this method. Duplicate reads resulting from PCR were identified and removed via Super Deduper (https://github.com/s4hts/HTStream). Then we used SeqPrep (https://github.com/jstjohn/SeqPrep) to simultaneously identify adapter contamination and overlapping paired reads, and then trim adapters and merge reads as necessary. We trimmed bases from both read ends via four bp sliding window to a minimum quality of 15 and then removed reads that were less than 30 bp long via Trimmomatic v2.X (Bolger, Lohse, & Usadel, 2014). Finally, we removed any remaining reads that were comprised of more than 50% of one nucleotide via remove_low_complex.py (distributed as part of the NF-Polish sequence polishing pipeline described in Irestedt et al. (2022)). We aligned cleaned sequencing reads to the Rufous-bellied Thrush (*T. rufiventris*) reference genome (ASM1318643v1) via BWA 0.7.17 *mem* (Li, 2013) and indexed mapped reads with Samtools 1.9 *index* (Danecek et al., 2021; Li et al., 2009). Following sequence cleaning and alignment we evaluated adapter contamination and sequence quality via FastQC. We used MapDamage 2.0 (Ginolhac, Rasmussen, Gilbert, Willerslev, & Orlando, 2011; Jónsson, Ginolhac, Schubert, Johnson, & Orlando, 2013) to estimate the frequency of C to T and G to A misincorporations that result from a transition to uracil during DNA degradation over time through hydrolysis. For each library we output sequencing metrics via Samtools 1.9 *stats*.

### 2.4. Analyses

We tested whether there were differences in DNA yield and mean DNA fragment size between different sources (toepad vs. skin clip) via paired-t tests. We evaluated whether DNA source, library preparation method, or an interaction between them resulted in differences in 1) endogenous DNA content, 2) sequencing efficiency, and 3) mean read length via two-way, repeated-measures ANOVAs. For any two-way ANOVA that resulted in a significant interaction, we performed a one-way, repeated measures ANOVA for each method to evaluate whether there were significant differences by DNA source. For any two-way ANOVA that resulted in a significant effect of either independent variable, we performed subsequent pairwise, paired-t tests between all library preparation methods. We accounted for multiple-testing in all post-hoc one-way and paired-t tests by adjusting p-values via the BH method (Benjamini & Hochberg, 1995).

We expected that samples with larger mean DNA fragment sizes would also have longer mean read lengths and as a result, greater endogenous DNA content and sequencing efficiency. To test this hypothesis while controlling for any effect of DNA source and library preparation method we defined two linear models for each of the following response variables: endogenous DNA content, sequencing efficiency, and read length. Each null model included the library preparation method and DNA source as fixed effects and sample as a random effect. The alternative model also included mean DNA fragment length as a fixed effect. To test whether mean DNA fragment length had a significant influence on each response variable we performed a likelihood ratio test of the null and alternative model.

Finally, we sought to qualitatively evaluate the effect of predigestion on replicate samples. To do so, we plotted the difference in each metric of interest between the replicate samples. All statistical analyses were completed in R v4.1.0 (R Core Team, 2021). ANOVAs and t-tests were conducted with the package *rstatix v0.7.0* (Kassambara, 2021), linear mixed models were built in *lme4* v1.1-27.1 (Bates, Mächler, Bolker, & Walker, 2015), and we used the suite of functions in *tidyverse* v1.3.1 (Wickham et al., 2019) for data parsing, manipulation, and visualization.

## RESULTS

### 3.1. DNA Yield and Size

All 60 DNA extractions were successful in terms of producing measurable amounts of DNA with an average of 589.1 nanograms (ng) per sample and a minimum of 48.6 ng in the skin clip control replicate from robin specimen 162188 (Table 1). All samples retained enough DNA through the DNA repair and cleanup to progress to library preparation by each of the three methods. In general, the DNA repair resulted in an upshift in the distribution of DNA fragment lengths (Figures S1A and S2A). There was no statistical difference in DNA yield and size between toepad and skin clips immediately following extraction or after the DNA repair and cleanup (Figure 2). However, the lack of statistical difference in DNA size is driven by the large variance in skin clips (Table 2). The mean size of DNA extracted from the toepad sample is greater than that of the skin clip sample for all but two specimens: robin specimen 83114 and thrush specimen 66823 (Table 1). For example, these two specimens bias the distribution of the post-repair skin clip mean size (post-repair M = 315.63, Mdn = 65.49, SD = 651.92) upwards, but not the toepad mean size (post-repair M = 83.94, Mdn = 83.94, SD = 7.13).

**Figure 2.**
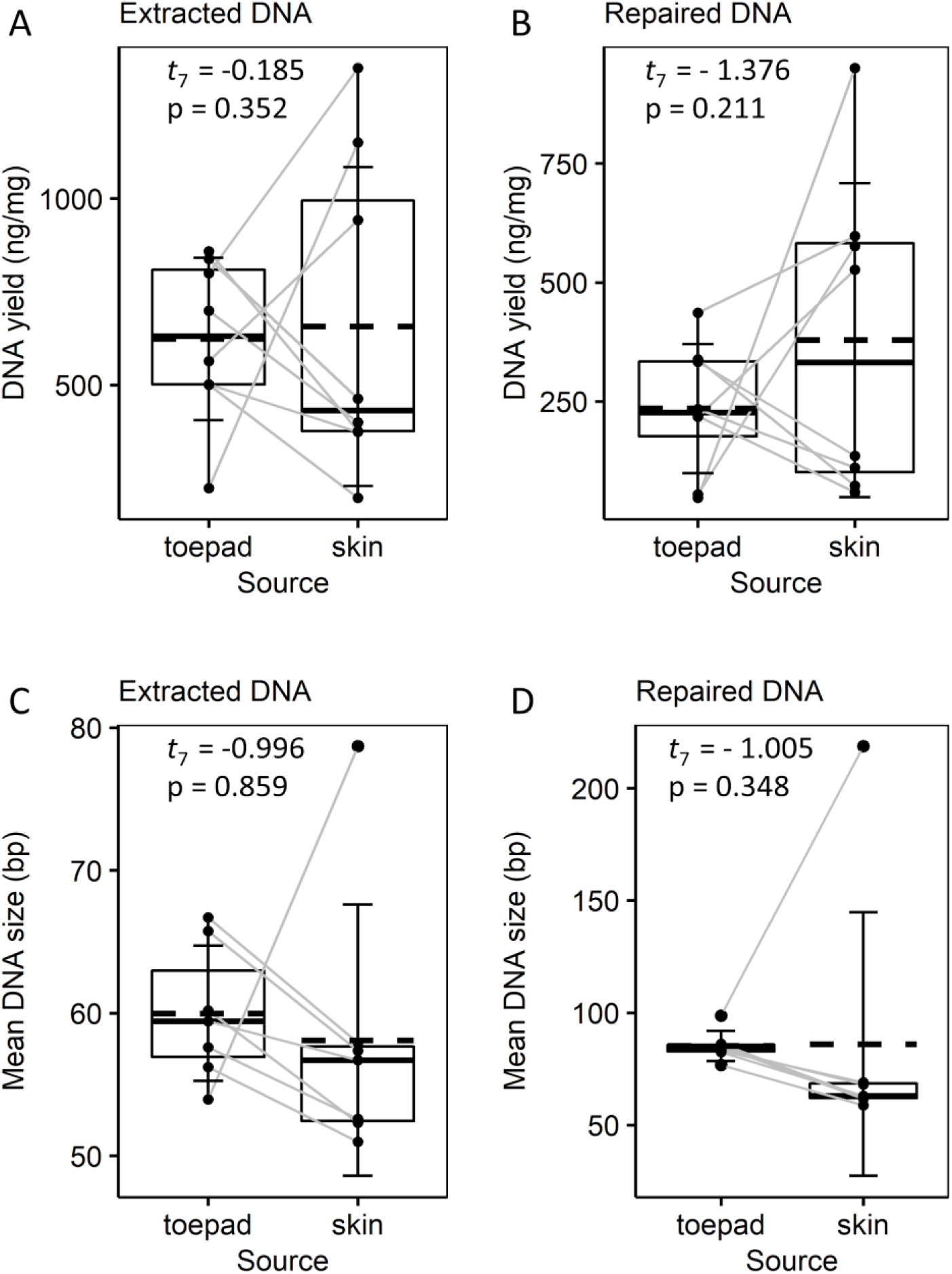
Source tissue impacts on DNA quantity and quality. For each sample, DNA yield (ng/mg) measured A) following DNA extraction and B) following DNA repair and cleanup is plotted by DNA source (toepad or skin clip). For each sample, mean DNA fragment size (bp) C) following DNA extraction and D) following DNA repair are plotted by DNA source. Samples from robin specimen 66823 are not plotted in C and D because the mean DNA size of its skin clip (Table 1) limits visualization of the variation in the size of the other samples. Gray lines connect toepad and skin clip data points from the same specimen. Summary statistics are also plotted by source: the mean and median are represented by dashed and solid lines, respectively, the upper and lower limits of the boxes represent the 75% and 25% quantiles, and the error bars represent the standard deviation.

**Table 2.**
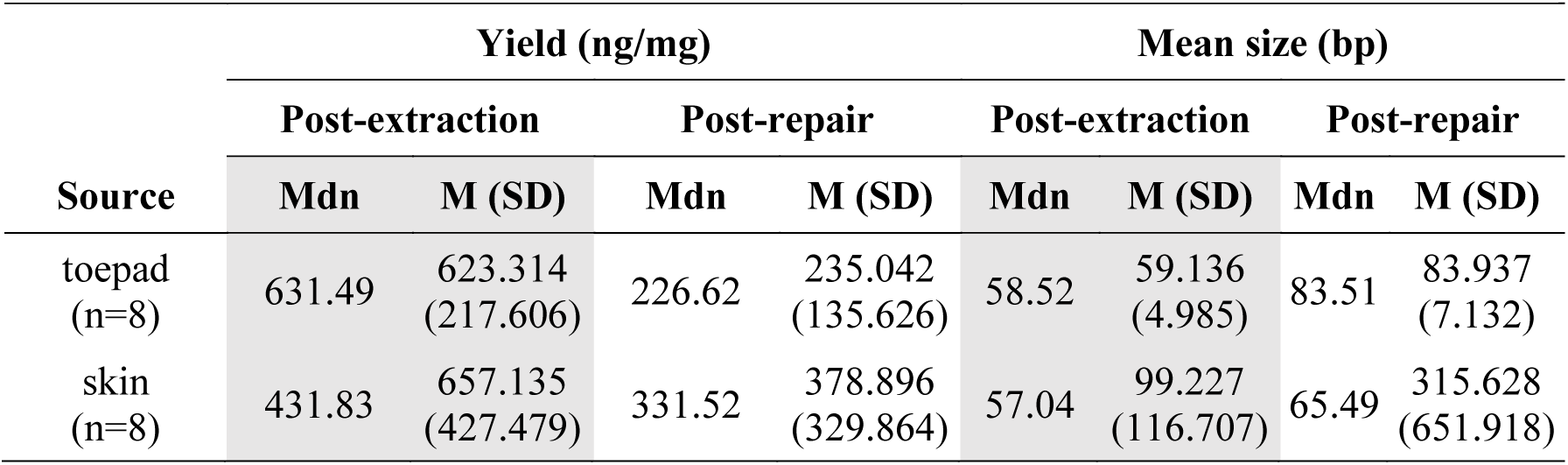
Summary (median, mean, standard deviation) of DNA yield and mean DNA size for predigested toepads and skin clips.

| Source | Yield (ng/mg) |  |  |  | Mean size (bp) |  |  |  |
| --- | --- | --- | --- | --- | --- | --- | --- | --- |
|  | Post-extraction |  | Post-repair |  | Post-extraction |  | Post-repair |  |
|  | Mdn | M (SD) | Mdn | M (SD) | Mdn | M (SD) | Mdn | M (SD) |
| toepad<br>(n=8) | 631.49 | 623.314<br>(217.606) | 226.62 | 235.042<br>(135.626) | 58.52 | 59.136<br>(4.985) | 83.51 | 83.937<br>(7.132) |
| skin<br>(n=8) | 431.83 | 657.135<br>(427.479) | 331.52 | 378.896<br>(329.864) | 57.04 | 99.227<br>(116.707) | 65.49 | 315.628<br>(651.918) |

### 3.2. Endogenous DNA content and sequencing efficiency

Sequencing returned a total of approximately 1.45 billion raw reads and, per library, an average of 11.23 million raw reads (SD = 1.84) and 9.88 million mapped reads (SD = 9.78) per library excluding non-predigested replicates. Detailed sequencing metrics for each library are reported in the supplementary material (Table S1).

There was a significant difference across library-preparation methods, but not DNA source in endogenous DNA content and sequencing efficiency with IDT outperforming SRSLY and KAPA in both metrics (Table 3). Similar to the results described above for DNA size, the toepad samples outperform the corresponding skin clip samples except for specimens 83114 and 66823 (Figure 3A, 3B) so we summarize the results by method and source (Table 4). The average endogenous DNA content of IDT toepad and skin clip libraries is 88.7% (SD = 0.014) and 81.2% (SD = 0.140) respectively, 0.06% and 1.4% greater than that of KAPA, and 2.4% and 1% greater than that of SRSLY. The average sequencing efficiency of IDT toepad and skin clip libraries is 35.9% (SD = 0.045) and 29.8% (SD = 0.245), respectively. In comparison to IDT, KAPA toepad and skin clip libraries are 11% and 1.8% less efficient, respectively, and SRSLY toepad and skin clip libraries are 7.8% and 1.8% less efficient. There was also a significant difference among methods in mean read length with IDT producing longer reads than KAPA and SRSLY with a significant interaction between method and DNA source (Table 3, Table 4, Figure 3C). IDT toepad libraries produced significantly longer reads than IDT skin clip libraries (Table 3, Table 4, Figure 3C).

**Table 3.**
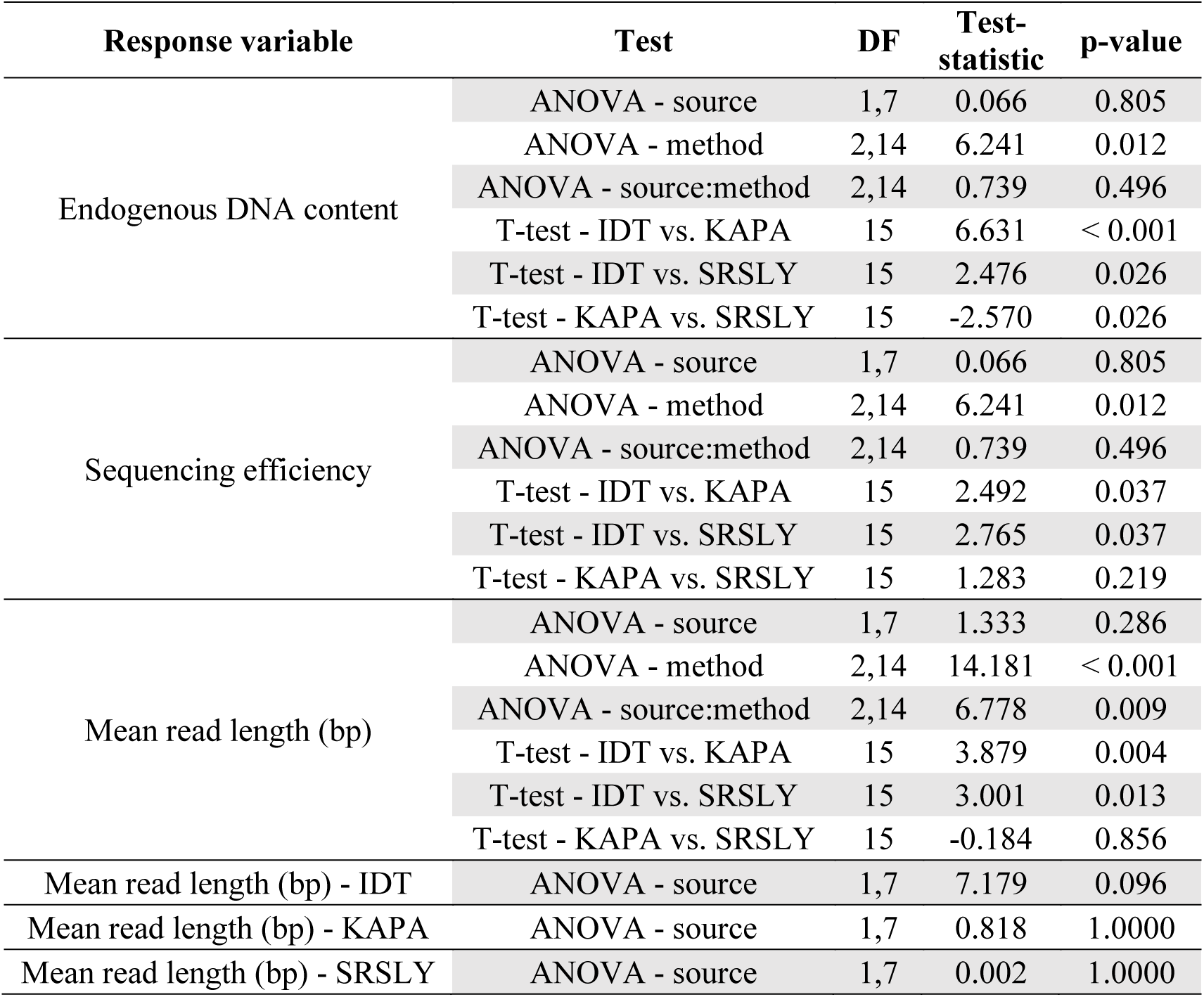
Summary of the statistical analyses of endogenous DNA content, sequencing efficiency, and mean read length, and their results. All the p-values presented for t-tests are corrected for multiple testing via the Benjamini and Hochberg (1995) method.

**Table 4.** Summary (median, mean, standard deviation) of sequencing metrics for predigested toepads and skin clips.

| Source | Mean read length (bp) |  |  |  |  |  | Endogenous DNA content |  |  |  |  |  | Sequencing efficiency |  |  |  |  |  |
| --- | --- | --- | --- | --- | --- | --- | --- | --- | --- | --- | --- | --- | --- | --- | --- | --- | --- | --- |
|  | IDT |  | KAPA |  | SRSLY |  | IDT |  | KAPA |  | SRSLY |  | IDT |  | KAPA |  | SRSLY |  |
|  | Mdn | M (SD) | Mdn | M (SD) | Mdn | M (SD) | Mdn | M (SD) | Mdn | M (SD) | Mdn | M (SD) | Mdn | M (SD) | Mdn | M (SD) | Mdn | M (SD) |
| toepad<br>(n=8) | 94.959 | 93.74<br>(5.598) | 78.428 | 78.998<br>(3.059) | 79.005 | 75.883<br>(9.771) | 0.892 | 0.887<br>(0.014) | 0.882 | 0.881<br>(0.009) | 0.865 | 0.863<br>(0.008) | 0.35 | 0.359<br>(0.045) | 0.245 | 0.249<br>(0.021) | 0.302 | 0.281<br>(0.072) |
| skin<br>(n=8) | 75.145 | 76.554<br>(14.799) | 60.717 | 71.669<br>(22.587) | 69.018 | 75.557<br>(19.097) | 0.846 | 0.812<br>(0.14) | 0.827 | 0.798<br>(0.14) | 0.823 | 0.802<br>(0.123) | 0.195 | 0.298<br>(0.245) | 0.151 | 0.236<br>(0.212) | 0.225 | 0.28<br>(0.179) |

**Figure 3.**
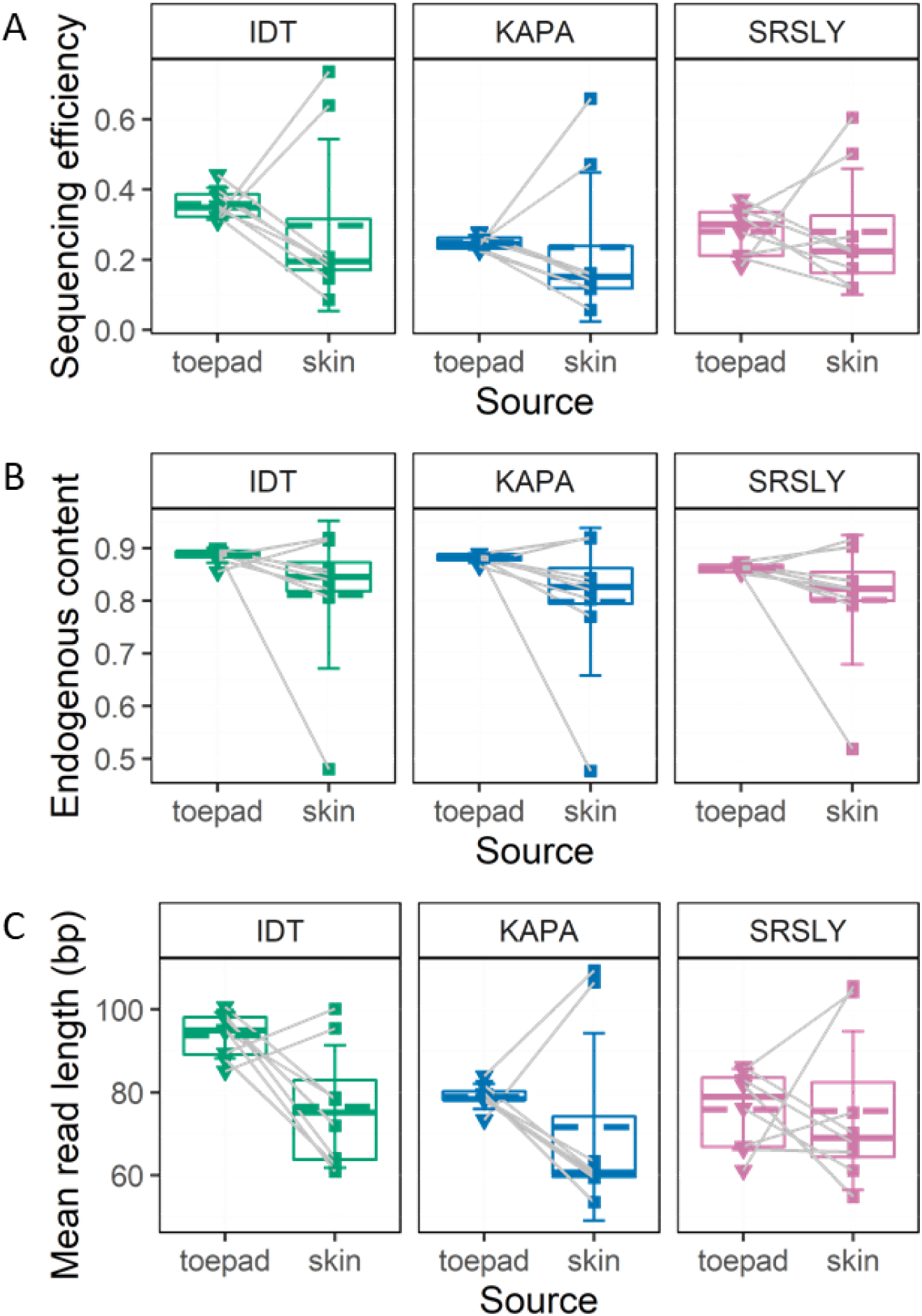
Association of sequencing library type and sequence data characteristics. For each library, A) sequencing efficiency, or the proportion of raw bases that uniquely mapped to the reference genome, B), endogenous DNA content, or the proportion of cleaned bases that uniquely mapped to the reference genome, and C) the mean read length (bp) is plotted by DNA source (toepad or skin clip). Gray lines connect toepad and skin clip data points from the same specimen. Summary statistics are also plotted by method and source: the mean and median are represented by dashed and solid lines, respectively, the upper and lower limits of the boxes represent the 75% and 25% quantiles, and the error bars represent the standard deviation.

The tests of the influence of input DNA fragment size on sequencing outcomes produced mixed results. Including DNA size significantly improved the fit of the linear models for endogenous DNA content (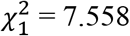, p = 0.006), sequencing efficiency (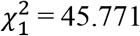, p < 0.001), and read length (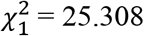, p < 0.001). However, in the linear models including DNA size, it was only a significant predictor of endogenous DNA content and read length based on confidence intervals of the coefficient estimate. In some cases, the trend for the relationship between DNA size and the dependent variable are contrary to our expectations (Figure S3).

There was evidence of contamination in two of the 20 libraries based on the proportion of base pair differences between the mapped reads and the reference genome and endogenous DNA content (Figure S4, Table 1). The skin clip library of thrush specimen 83109 exhibited approximately 1% more differences from the reference genome than all other samples (Table 1, Figure S4) in addition to a comparatively low proportion of endogenous DNA content across preparation methods (IDT = 48.0%, KAPA = 47.7%, SRSLY = 51.9%). The replicate skin clip library of robin specimen 162188 that was not subjected to predigestion also had a low proportion of endogenous DNA content across preparation methods (IDT = 17.4%, KAPA = 33.4%, SRSLY = 41.5%), but it was similar to all other libraries in the proportion of mapped bases that differed from the reference (Figure S4). There was no indication of contamination in the high performing skin clips samples from specimens 83114 and 66823 (Table 1, Figure S4) that biased the skin clip averages of most metrics upward.

The influence of sample predigestion was unclear (Table 1, Figure S5). The differences in DNA yield, mean DNA size, and endogenous DNA content were inconsistent between the control and predigested replicates from robin 162188 and thrush 83114. The difference between the replicates in sequencing efficiency and mean read length were marginal.

## 4. Discussion

### 4.1. WGS of hDNA from 100-year-old bird study skins

We have demonstrated via shallow sequencing of 60 hDNA libraries that ssDNA library preparation methods outperform dsDNA methods in sequencing efficiency and, to a lesser extent, in returning endogenous DNA content from WGS ~100-year-old bird specimens. In contrast to our predictions, the IDT multi-tube, ssDNA method outperformed the Claret Biosciences single-tube, ssDNA method and we discuss possible explanations below. We also confirm previous research suggesting that toepads provide consistently larger DNA fragments and demonstrate that hDNA from toepads rather than skin clips is a better source for WGS. We show that, though skin clips may sometimes outperform toepads for a given specimen, toepads have less variance and therefore less unexpected sequencing outcomes. Altogether, we’ve shown that toepads are a better source of DNA and ssDNA library preparations are a better method for collecting WGS from 100-year old bird specimens. Below we elaborate further on the nuances of our findings and conclude with broader implications for WGS of historical DNA from museum specimens in natural history collections.

### 4.2. Library preparation method and DNA source influenced sequencing

Library preparation method influenced all key metrics; but contrary to our predictions, the SRSLY ssDNA, single-tube method did not outperform the other two methods. Instead, the IDT ssDNA, multi-tube method resulted in greater endogenous DNA content, sequencing efficiency, and read lengths than either of the other methods (Figure 3, Table 3). The margin of difference between IDT and the other methods was much greater for sequencing efficiency and average read length than endogenous DNA content (Table 4), suggesting that IDT produced more complex libraries. The only other study to compare ssDNA and dsDNA methods for shotgun sequencing of historical specimens prepared libraries from beetle specimens of various ages using the same ssDNA, multi-tube method that we implemented and a different dsDNA, single-tube method. Those results showed no difference in endogenous DNA content between methods, and that ssDNA libraries maintained more sequencing reads through quality filtering and trimming (Sproul & Maddison, 2017). We find this consistent with our results and suspect that controlling for taxonomy and specimen age in our experimental design facilitated detecting the small difference that library preparation method made in endogenous DNA content. That IDT outperformed SRSLY makes some sense given that SRSLY was originally developed for cell-free DNA which averages 30 bp long (Troll et al., 2019); though, the commercial kit provides several versions of the protocol optimized for different purposes, and we implemented the version for moderate length DNA inserts, less than 200 bp. That protocol for moderate length DNA inserts includes two-sided SPRI cleanups following adapter ligation and indexing PCR, as compared to the IDT ssDNA method which uses single-sided cleanups. We suspect that the two-sided cleanups in SRSLY limited conversion of DNA molecules on the larger end of the DNA size distribution into library molecules (Figures S1, S2). This is consistent with SRSLY producing shorter average read lengths than IDT. Still, SRSLY outperformed the KAPA dsDNA method overall in sequencing efficiency and outperformed IDT in sequencing efficiency for skin clips in a few specimens (Figure 3A). It is possible that SRSLY may be the better method for samples that are more degraded as a result of age or DNA source. More recently another ssDNA single-tube method that builds upon SRSLY was developed specifically for ancient DNA samples (Kapp et al., 2021); though the nonproprietary status limits ease of implementation. It may be worthwhile to optimize the SRSLY clean up steps to maximize conversion of the largest DNA fragments as the IDT method costs 1.89× more than SRSLY per reaction.

Most statistical tests of the effect of DNA source on our metrics of interest were not significant, though toepads seem to perform better for WGS when considering the influence of the large variance in the skin clip metrics. Toepads had much smaller variance in DNA yield, DNA size, endogenous DNA content, sequencing efficiency, and average read length than skin clips. For six of the eight specimens, toepads clearly provided greater endogenous DNA content, sequencing efficiency, and average read length than the corresponding skin clip. In general, this result is reflected by the mean, and more so the median values, of these metrics for toepads as compared to skin clips (Table 2, Table 4). That phenol chloroform extraction of toepads does not yield more DNA but does result in longer DNA fragments than that of skin clips supports previous research (Tsai et al., 2020). Our results are also in line with previous studies based on five fluid-preserved garter snake specimens (Zacho et al., 2021) and three dried mammal skins from different species (McDonough et al., 2018) that showed that different DNA sources have differences in endogenous DNA content. It is possible that hDNA sampled from bird specimen toepads produces better WGS data than that of skin clips because of their different structural makeups. The keratinized, scaly skin of bird feet may provide a better environmental barrier to water—which promotes DNA degradation via hydrolysis and also overall tissue degradation— and to invading microbes that would increase exogenous DNA. Toepad cells may also have lower innate water content due to desiccation during keratinization (Bengtsson et al., 2012). The role of the keratin structure in maintaining better DNA for WGS is supported by the work of McDonough and colleagues (2018) which showed that of bone, skin, and claw samples from dried mammal specimens, claw samples had the highest or near highest proportion of endogenous DNA and in a qPCR analysis, the highest copy number of nuclear genomic markers.

### 4.3. Predigestion, high performing samples, and potential contamination

Interpretation of the effect of predigestion on reducing exogenous DNA content and increasing sequencing efficiency is limited by the small sample size for which we sequenced replicate predigested and control samples. The lack of clear signal of predigestion effect in DNA yield and size is unsurprising given the variation across all samples (Figure 2). Moreover, the lack of any potential signal is unsurprising given that of the eight samples included as replicates to evaluate predigestion, three were exceptions to the general trends identified by the larger study. Both skin clips from thrush 83114 were among the few skin clip samples that performed uncharacteristically better than all other samples and the control skin clip replicate from robin from 162188 resulted in the low endogenous DNA content of all libraries (Table 1) indicating high levels of exogenous DNA or contamination. Notably, the predigested skin clip replicate sample from robin 162188 did not show signs of contamination suggesting that predigestion may have reduced exogenous DNA in only two minutes of predigestion time. In contrast, the skin clip from thrush specimen 83109 was one of four samples with the shortest, one-minute predigestion and also showed a clearer signal of contamination based on low endogenous DNA content and a larger genetic distance from the reference than other samples (Table 1, Figure S4). Finally, it is likely that predigestion reduced total DNA yield, though enough DNA remained for all 20 samples to prepare three successfully sequenced libraries. Altogether we suggest this preliminary investigation is a promising avenue to maximize endogenous hDNA from museum bird specimens for WGS and warrants further research.

The primary source of the larger variance in most metrics for skin clips were two specimens for which the skin clips not only outperformed the corresponding toepad from the same specimen but all other samples. Robin 66823 and thrush 83114 returned the first and second largest DNA yield, DNA size, endogenous DNA content, average read length, and highest sequencing efficiency. Notably, thrush 83114 was one of the two specimens included in the predigestion study, and both the predigested and non-predigested skin clips were high performing samples and had consistent values across metrics. There are multiple explanations for why the samples outperformed all others, the first being contamination. However, neither of these samples show clear signs of contamination like the lower endogenous DNA content or large genetic distance from the reference genome mentioned for the skin clip from thrush 83109. The only explanation of contamination we consider plausible is contamination by modern DNA from the same or a closely related species, perhaps from a more recently collected specimen in the same drawer. This possibility highlights the need for stringent sampling procedures during sample collection from museum specimens for hDNA purposes. Importantly, a population genomics study involving deeper sequencing of these samples would allow assembly of mitochondrial genomes that would enable identification of multiple individuals within one sequencing library. Another possible explanation for the higher performance of these skin clips is that these specimens received different treatments at the time of collection than the other specimens in the study. At the time of collection it was common practice to treat bird (and mammal) skins with arsenic-containing solutions for tanning as well as protection from pests (Marte, Péquignot, & Von Endt, 2006), and arsenic has been demonstrated as a DNA polymerase inhibitor (Töpfer et al., 2011). It is possible that the specimens with high performing skin clips were accidentally skipped in some treatment that ultimately promoted DNA damage in the other skins. The last and, in our opinion, most likely possibility is that this variation represents real variation in DNA quality between specimens. Such large variations are not uncommon when working with hDNA (e.g. McDonough et al., 2018) and ancient DNA (e.g. Wales et al., 2015) and highlights the value of identifying methods and DNA sources that can consistently return an expected amount of WGS data like we have found for toepads and the IDT ssDNA, single-tube method.

## 5. Conclusion

We have shown that for 100-year old museum bird study skin specimens, of those combinations we tested, the combination of toepads and ssDNA library preparation, in this case the IDT xGen™ ssDNA & Low-Input DNA Library Prep Kit, provide the best WGS data. Our results regarding toepads, in combination with previous studies of endogenous DNA content in other taxa, can be reasonably extended beyond birds to suggest that keratinous sources of hDNA may be the best source for WGS and should motivate additional investigations of this hypothesis. We also have shown that when comparing WGS from ssDNA and dsDNA methods, ssDNA methods provide a larger increase in sequencing efficiency than endogenous DNA content, suggesting that they successfully convert more hDNA molecules into sequenceable library molecules and likely lead to more complex libraries better suited for WGS at the depth required for population genomic studies. Further study of the impact of library preparation method on sequencing efficiency that controls for variation among specimens and also evaluates the role of age of the specimen is necessary to identify the threshold at which an ssDNA method is or is not warranted. Also, it may be worthwhile to attempt to further optimize SRSLY cleanups to minimize bias against converting larger fragment hDNA molecules into library molecules as SRSLY is currently ~89% less expensive per reaction than the IDT method. Finally, our inclusion of a predigestion step to reduce external exogenous DNA did not yield straightforward results, though it did provide some evidence that it limited contamination in one of four samples for which we made a direct comparison. Importantly we showed that a brief (less than 15 minute), gentle (37C° incubation) predigestion step does not preclude successful library construction, and thus we will cautiously include this step in our own protocols moving forward. Altogether this study identifies how to maximize WGS data collected from 100-year old bird specimens and provides some general insights on how to increase the quality and quantity of WGS data recovered from hDNA of museum specimens overall.

## Acknowledgments

This work was supported by NSF Grant No. 1953796 to JDM and BDM. We would like to thank Dr. Manaasa Raghavan for providing access to the ancient DNA facility at the University of Chicago and the Raghavan Genoscape Lab, especially Dr. Constanza de la Fuentes, for facilitating our pre-PCR lab work. Mohamed Fokar at the TTU Center for Biotechnology & Genomics provided sequencing support. The TTU Center for Biotechnology & Genomics acquisition of the NovaSeq6000 was supported by NIH grant 1S10OD025115-01. We acknowledge the High Performance Computing Center (HPCC) at Texas Tech University (http://www.hpcc.ttu.edu) and the Pritzker DNA Lab at Field Museum of Natural History for providing computational and molecular resources, respectively, that have contributed to the research results reported within this paper.

## Data Accessibility and Benefit-Sharing

Raw sequence reads will be deposited in NCBI Sequence Read Archive (SRA) (Settlecowski et al. 2022). Code used in data analysis is available at: github.com/amiesett/WGS-hDNA-birds. Benefit sharing: We are sharing all research benefits of this project by providing raw data, bioinformatic scripts, and results in public databases. No monetary benefits are expected from these research results.

## Author Contributions

All authors contributed to project design and manuscript completion. JDM and BDM acquired funding. AES performed lab work and data analysis, with support from JDM, and wrote the first draft of the manuscript.

