## Supplemental figures for "Library preparation method and DNA source influence endogenous DNA recovery from 100-year-old avian museum specimens"

**Running title:** WGS of historical avian museum specimens

Amie E. Settlecowski [†1]

Ben D. Marks[1]

Joseph D. Manthey [†2]

[†]corresponding author

[1]Bird Collection, Field Museum of Natural History, Chicago, Illinois, USA

[2]Department of Biological Sciences, Texas Tech University, Lubbock, Texas, USA

### Supplemental Figures

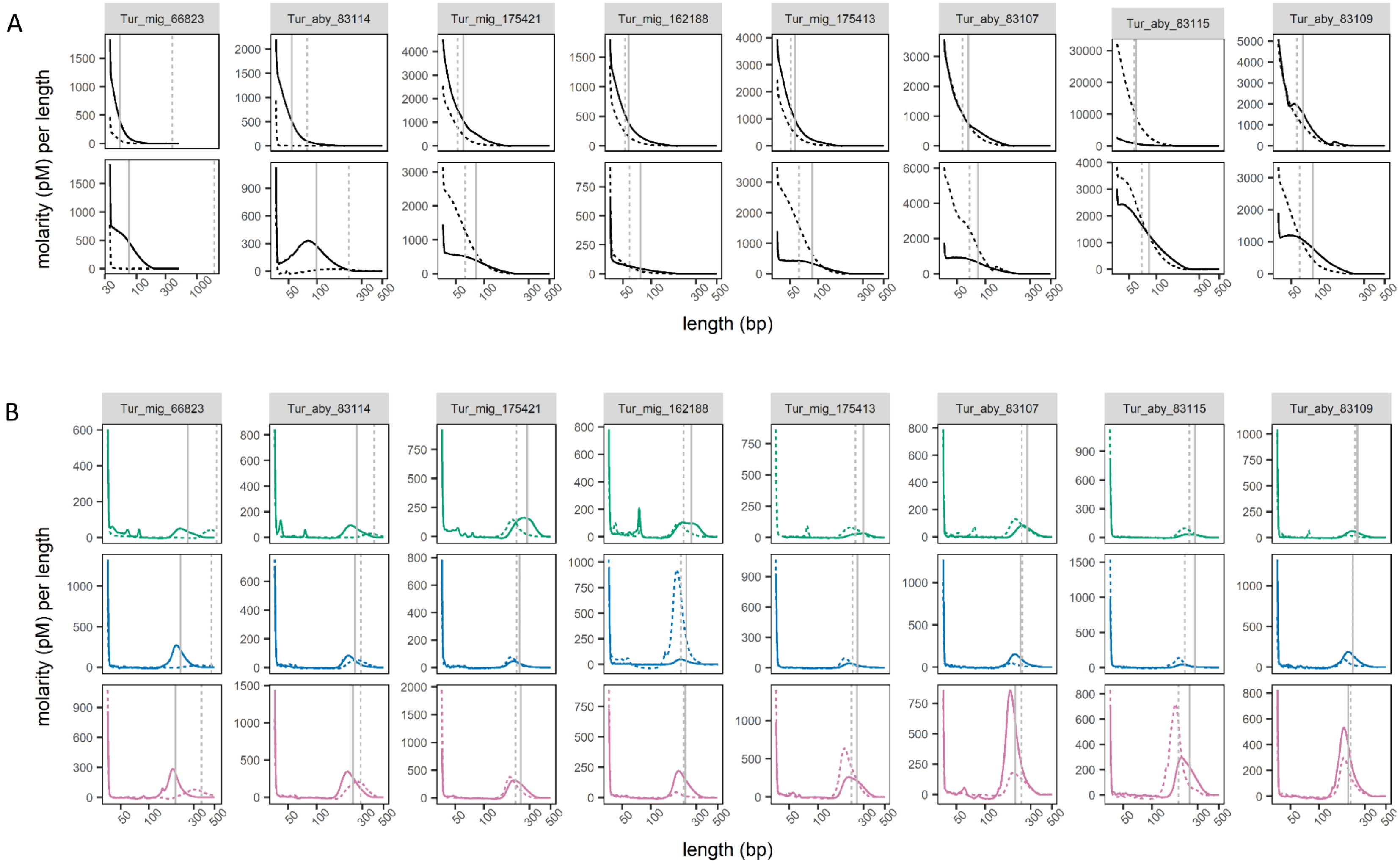


**Figure S1.** DNA fragment length distributions. Outputs from Bioanalyzer are plotted as molarity per length. Solid and dashed lines represent toepad and skin clip samples, respectively. Gray vertical lines represent the mean fragment lengths. A) Top and bottom rows are from DNA extractions and repaired DNA extractions, respectively. B) The rows correspond to IDT (green), KAPA (blue), and SRSLY (pink) libraries for the specified specimen.

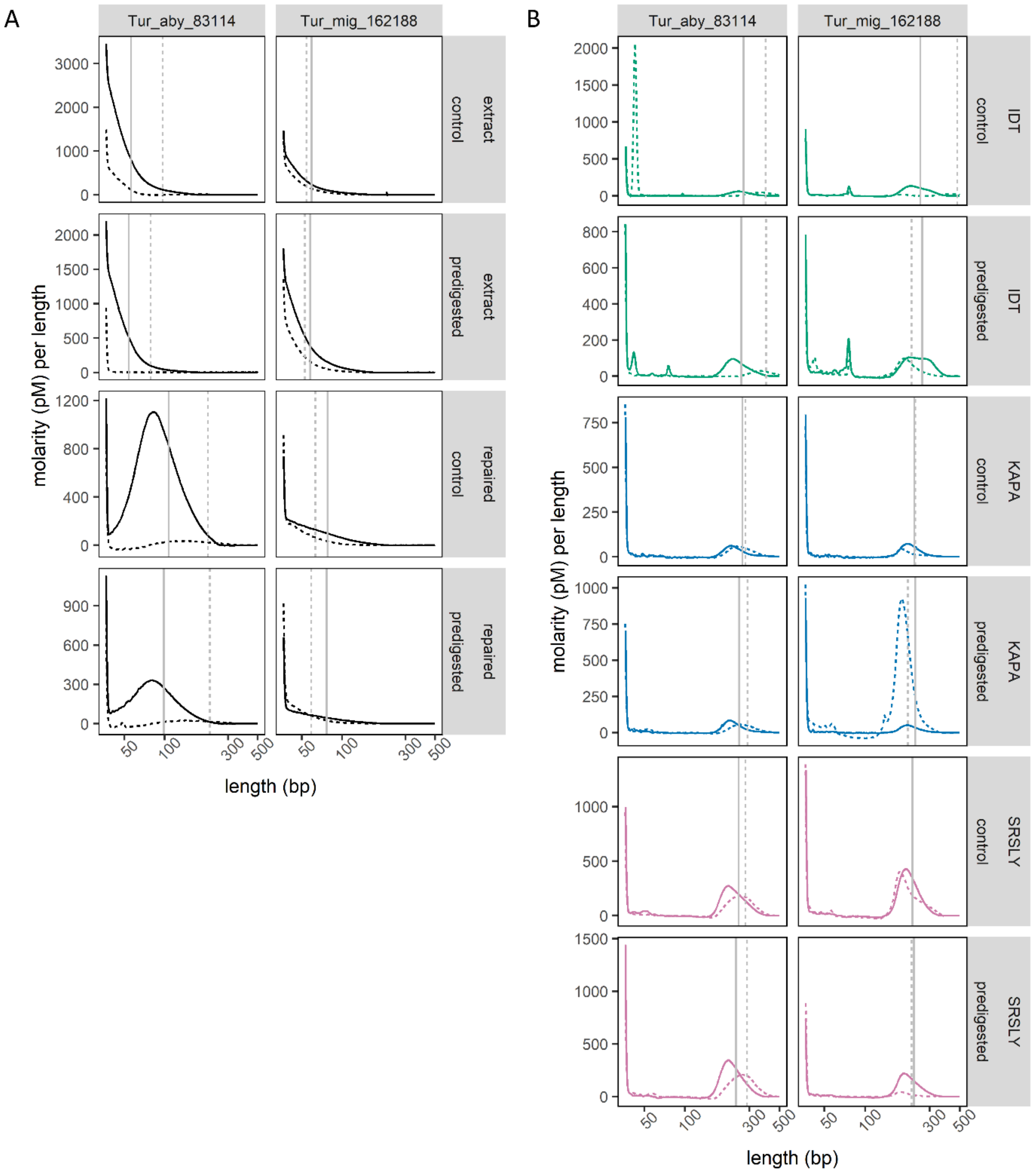


**Figure S2.** DNA fragment length distributions of replicate control and predigestion samples. Outputs from bioanalyzer are plotted as molarity per length. Solid and dashed lines represent toepad and skin clip samples, respectively. Gray vertical lines represent the mean fragment lengths. A) Plots show outputs from DNA extractions and repaired DNA extractions. B) Plots show outputs from libraries.

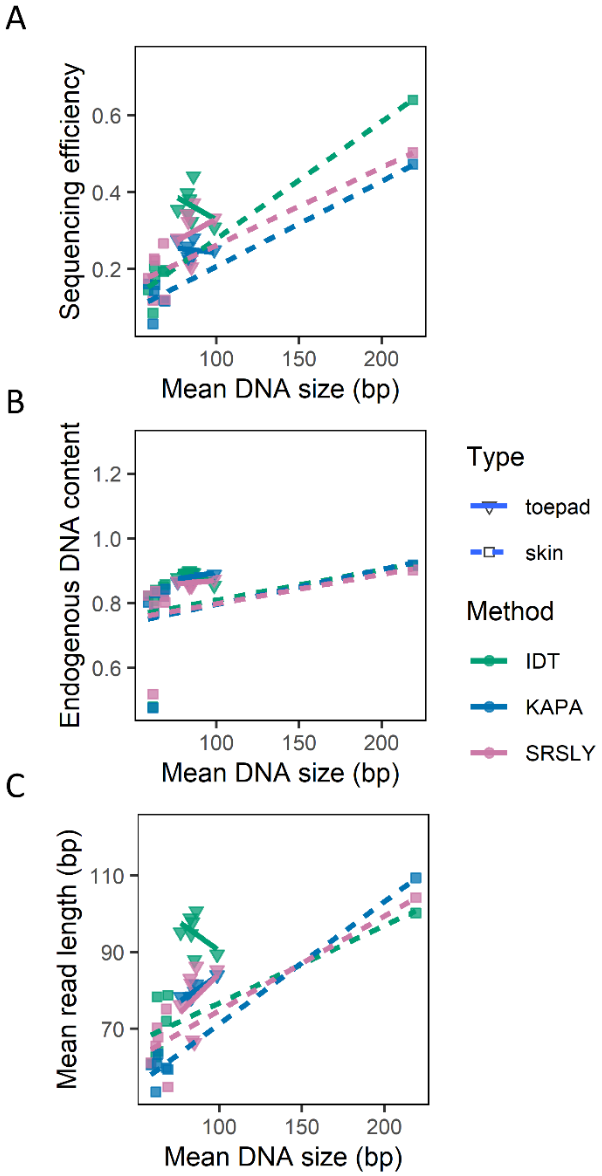


**Figure S3.** Association between DNA input size and sequence data characteristics. For each library, A) sequencing efficiency, B) endogenous DNA content, and C) read length (bp) is plotted against mean DNA size (bp). For each combination of library preparation method and DNA source, the relationship between each metric and DNA size is presented by linear regression. Libraries prepared from robin specimen 66823 are not shown because the mean DNA size of its skin clip (Table 1) limits visualization of the relationship in the other samples.

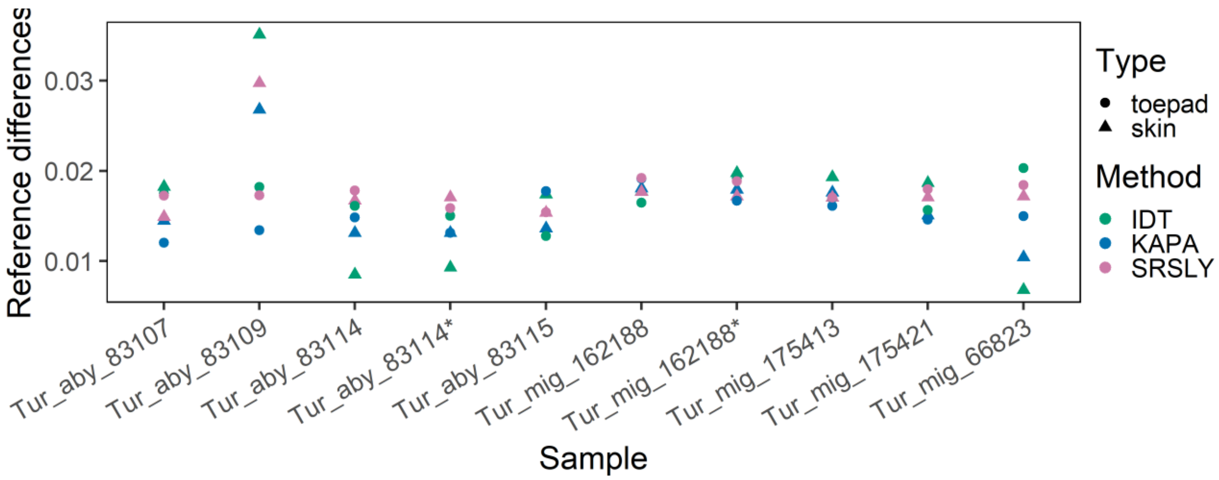


**Figure S4.** Genetic distance of mapped sequences to the reference genome. Genetic distance measured as the proportion of bases mapped to the reference genome that differ from the reference sequence. The two replicate samples that were not predigested are indicated by *.

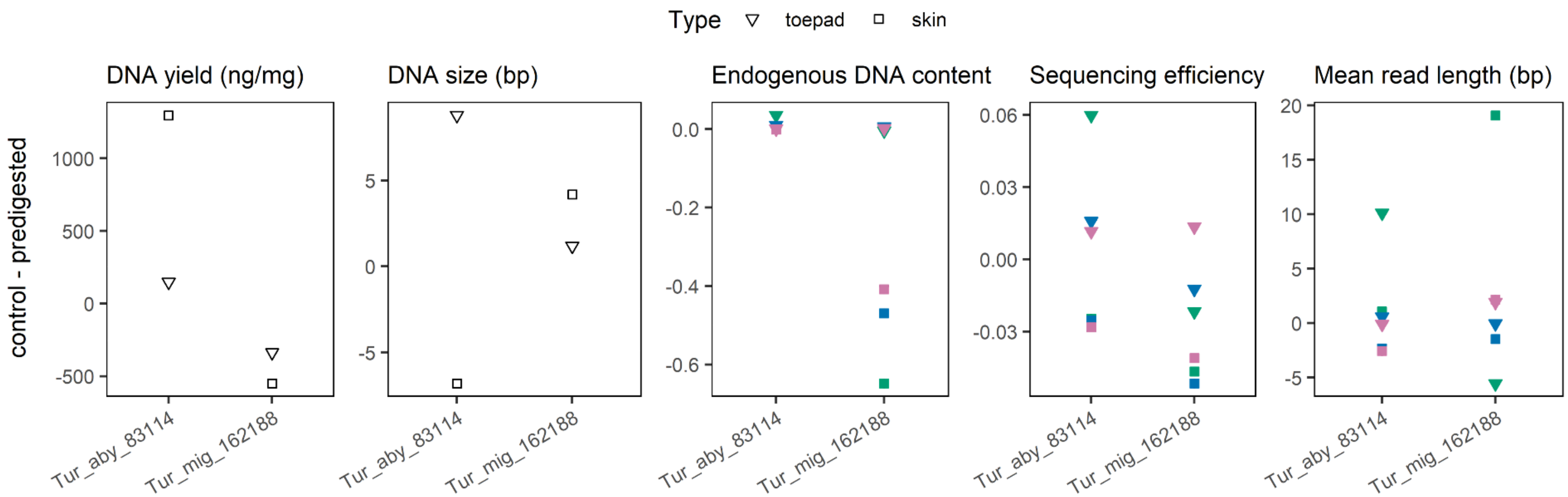


**Figure S5.** Predigestion impacts. The difference between two control and predigested replicate toepad and skin samples are plotted by specimen for each metric of interest. Green, blue, and pink correspond to the IDT, KAPA, and SRSLY libraries, respectively.
